# BLM-dependent Break-Induced Replication handles DSBs in transcribed chromatin upon impaired RNA:DNA hybrids dissolution

**DOI:** 10.1101/2020.05.13.093112

**Authors:** S Cohen, A Guenolé, A Marnef, T Clouaire, N Puget, V Rocher, C Arnould, M Aguirrebengoa, M Genais, D Vernekar, R Mourad, V Borde, G Legube

**Author notes:** These authors contributed equally (S Cohen, A Guenolé).

## Abstract

Transcriptionally active loci are particularly prone to breakage and mounting evidence suggest that DNA Double-Strand Breaks arising in genes are handled by a dedicated repair pathway, Transcription-Coupled DSB Repair (TC-DSBR), that entails R-loops accumulation and dissolution. Here, we uncovered a critical function of the Bloom RecQ DNA helicase (BLM) in TC-DSBR in human cells. BLM is recruited in a transcription dependent-manner at DSBs where it fosters resection, RAD51 binding and accurate Homologous Recombination repair. However, in a R-loop dissolution-deficient background BLM switches from promoting Homologous Recombination to promoting Break-Induced Replication (BIR), which strongly impairs cell viability. Altogether our work unveils a role for BLM in BIR at DSBs in active chromatin, and highlights the toxic potential of RNA:DNA hybrids that accumulate at these transcription-associated DSBs.

## Introduction

DNA Double-Strand breaks (DSBs) are harmful lesions that occur on the genome following exposure to various environmental sources such as radiation or chemotherapy, but that also arise on a regular basis due to cell metabolic activity such as during replication or the release of topological stress. Recently, genome-wide sequencing based analyses unveiled that endogenous DSBs primarily occur in genomic loci prone to form secondary structures, such as G-quadruplexes (G4), transcribed, or CTCF-bound loci (for instance (Canela et al., 2017, 2019; Crosetto et al., 2013; Dellino et al., 2019; Gothe et al., 2019; Lensing et al., 2016) reviewed in (Marnef et al., 2017; Puget et al., 2019)).

While DSB repair pathways, including non-homologous End Joining (NHEJ) and Homologous Recombination (HR), have been well characterized (reviewed in (Scully et al., 2019)), recent evidence suggests that repairing DSB in RNA Pol II-transcribed loci requires additional mechanisms, collectively referred to as Transcription-coupled DSB repair (TC-DSBR) (Marnef et al., 2017; Tan and Huen, 2020). In post-replicative cells, transcribed loci are specifically channelled to a specific homologous recombination repair pathway (Aymard et al., 2014) while in G1 these damages tend to persist and cluster (Aymard et al., 2017) (reviewed in (Marnef et al., 2017; Puget et al., 2019)). The G2 specific arm of TC-DSBR, also known as TAHRR (for Transcription Associated Homologous Recombination Repair) (Yasuhara et al., 2018), entails R-loops accumulation and resolution in a Senataxin (SETX), XPG, DDX1, and EXOSC10 dependent manner (Bader and Bushell, 2020; Cohen et al., 2018; Domingo-Prim et al., 2019; Li et al., 2016; Yasuhara et al., 2018) (reviewed in (Puget et al., 2019)). The mechanism(s) that accounts for RNA:DNA hybrids accumulation is still under investigation, but has been proposed to ground on either transcriptional arrest induced at damaged active genes (Kakarougkas et al., 2014; Meisenberg et al., 2019; Shanbhag et al., 2010) (reviewed in (Puget et al., 2019)) or on DNA end-mediated *de novo* transcription (D’Alessandro et al., 2018) and requires the miRNA processing enzyme Drosha (Lu et al., 2018) as well as resection (D’Alessandro et al., 2018; Yasuhara et al., 2018) (for review (Jimeno et al., 2019; Puget et al., 2019)). While RNA:DNA hybrids further contribute in recruiting RAD52 (Yasuhara et al., 2018) and BRCA2 (D’Alessandro et al., 2018), their resolution is mandatory to ensure completion of homologous recombination (Cohen et al., 2018; Domingo-Prim et al., 2019; Li et al., 2016; Yasuhara et al., 2018). This is highlighted by the strong survival defect described upon depletion of the SETX RNA:DNA hybrids helicase, which is only observed when DSBs are induced in active loci, and not randomly across the genome (Cohen et al., 2018).

The Bloom Syndrome helicase (BLM) is a 3’ to 5’ DNA helicase, mutated in the Bloom Syndrome, which is a genetic disorder associated with an increased risk of cancer, sun-induced chronic erythema, impaired fertility and immune deficiency. BLM displays pleiotropic functions in response to DSBs. First, it contributes to HR including in the dissolution of double Holliday junctions (Cejka and Kowalczykowski, 2010; Cejka et al., 2010a; Wu and Hickson, 2003), in heteroduplexes rejection (Mendez-Dorantes et al., 2020) and in promoting long-range resection (Cejka et al., 2010b; Gravel et al., 2008; Mendez-Dorantes et al., 2020; Nimonkar et al., 2011; Pinto et al., 2016; Soniat et al., 2019; Sturzenegger et al., 2014; Xue et al., 2019) (for review (Croteau et al., 2014; Marini et al., 2019)). Second, BLM also displays anti-recombinogenic properties, since (i) it promotes 53BP1 (an anti-resection factor) foci assembly and acts together with 53BP1/RIF1 to protect against CtIP-dependent long-range deletions (Grabarz et al., 2013; Tripathi et al., 2007, 2008), (ii) it disrupts RAD51 filament assembled on ssDNA and inhibit D-loop formation (Bugreev et al., 2007) and (iii) its depletion rescues RAD51 loading in an HR-impaired, BRCA1Δ11 mutant, background (Patel et al., 2017). Finally, BLM (or its yeast counterpart sgs1) was also proposed to contribute to Break-Induced Replication (BIR) (Mehta et al., 2017) and BIR-like pathways such as during Alternative Lengthening of Telomeres (ALT) that elongates telomeres in telomerase negative cells (Dilley et al., 2016; Lu et al., 2019; Min et al., 2019; Roumelioti et al., 2016; Silva et al., 2019; Sobinoff et al., 2017; Zhang et al., 2019).

Of interest, previous work revealed that BLM displays the ability to bind and unwind G-quadruplexes (G4) (Chatterjee et al., 2014; Lansdorp and van Wietmarschen, 2019; Sun et al., 1998; Tippana et al., 2016; Wu et al., 2015). Given the high prevalence of G4 at promoters and transcribed loci (Hänsel-Hertsch et al., 2016) this raises the interesting possibility that BLM may be a *bona fide* component of TC-DSBR which activity may be particularly relevant at transcribed, G4-forming, damaged loci. In agreement, Strand-seq experiments revealed that sister chromatid exchange (SCEs) breakpoints observed in BLM-deficient cells are biased toward G4 motifs, especially those present in transcribed genes (van Wietmarschen et al., 2018).

Here, in order to investigate the function of BLM during TC-DSBR we performed the first genome-wide distribution of BLM, at a high resolution, and at multiple DSBs induced simultaneously in transcribed and un-transcribed loci in human cells. We found that the recruitment of BLM is biased towards damaged transcribed and G4-prone loci and depends on transcriptional activity. At these breaks, BLM contributes to resection and RAD51 recruitment, ensuring faithful repair. However, we also uncovered a second role for BLM in promoting Break-Induced Replication (BIR) at DSBs in damaged genes, in the context of excessive RNA:DNA hybrids accumulation, upon SETX depletion. In agreement, genomic analyses in pancreatic cancer patient samples revealed that expression of SETX and BLM relate to the occurrence of BIR-associated mutational signatures. Altogether our data suggest that RNA:DNA hybrids which accumulate at DSBs in transcriptionally active loci are highly toxic intermediates that foster the use of a BLM-dependent cytotoxic and highly mutagenic BIR-like pathway.

## Results

### BLM is recruited at DSB induced in transcribed and G4-enriched loci

In order to get insights into the function of BLM during DSB repair, we analyzed its genome-wide distribution at a high resolution by ChIP-seq using our previously described DIvA system (DSB Inducible by AsiSI) (Iacovoni et al., 2010; Massip et al., 2010). This cell line expresses a restriction enzyme (AsiSI) fused to the ligand binding domain of the estrogen receptor. Following a treatment with 4-hydroxytamoxifen (4-OHT), the fusion enzyme is relocated in the nucleus, where it induces, homogeneously in the cell population, multiple DSBs at annotated positions located both in transcribed (promoters, genes bodies) and untranscribed loci (silent genes, intergenic) (Clouaire et al., 2018). ChIP-seq analyses post damage indicated that BLM is recruited at the vicinity of DSBs (Fig. 1A-B). Average BLM profile and heatmaps at all DSBs showed that BLM spreads on roughly 5-10 kb around DSBs and peaks at a position where γH2AX is depleted (Fig. 1C-D). These data therefore indicate that the BLM helicase is distributed on restricted domains around DSBs unlike the megabase-wide spreading observed for γH2AX.

**Figure 1.**
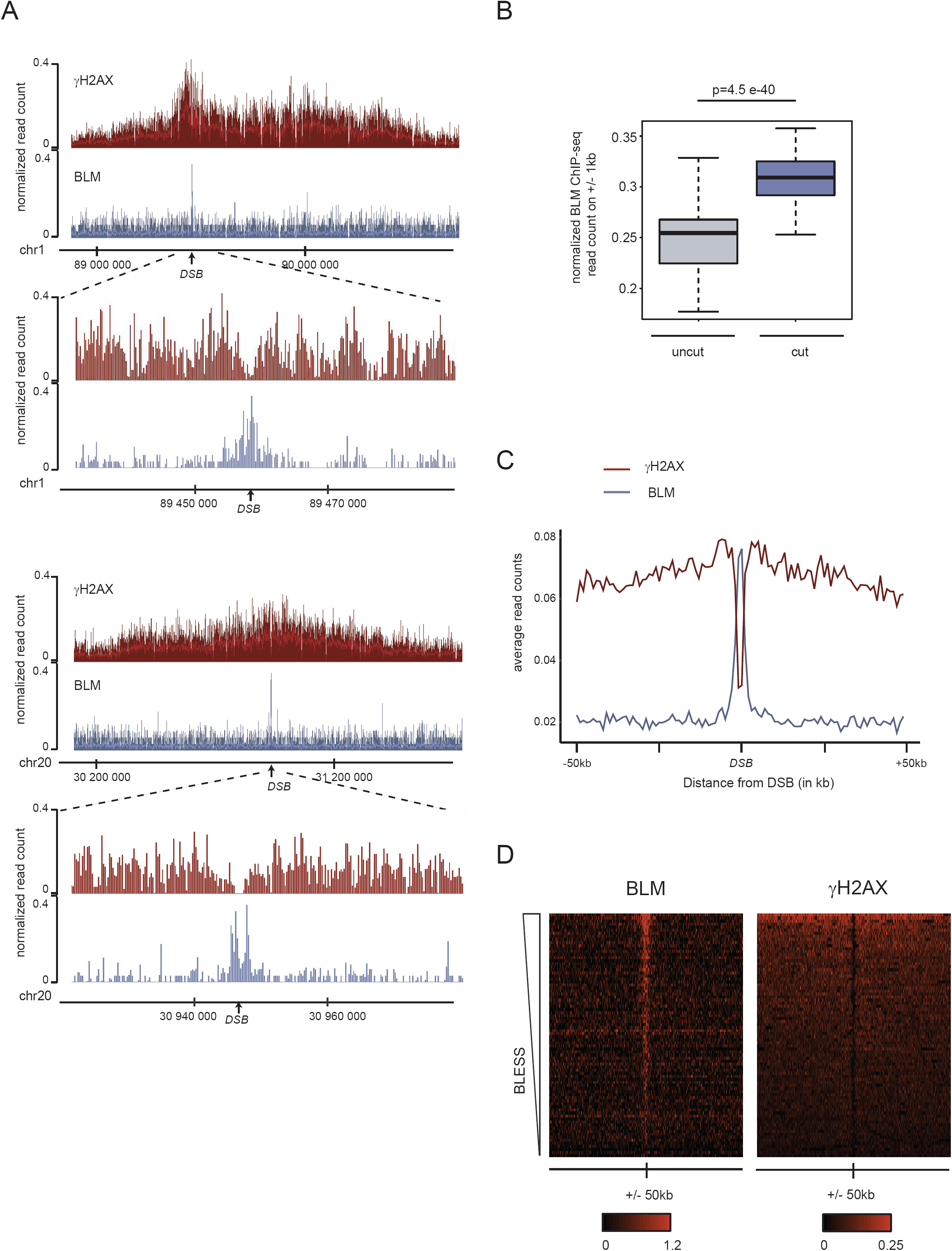
BLM recruitment and distribution at DSBs. (A) Genome browser screenshot representing γH2AX and BLM ChIP-seq signal at two DSBs (chr1: 89458586 (upper panel) and chr20: 30946312 (lower panel)). Magnifications are also showed. DSBs are indicated by arrows. (B) Boxplot representing BLM normalized ChIP-seq count, on a 2 kb window around the eighty-best induced DSBs (blue) and the 1,131 uncut AsiSI sites on the genome (grey). *P* values were calculated using two-sample Wilcoxon tests. (C) Average γH2AX (red) and BLM (blue) ChIP-seq read count profiles for the eighty-best induced DSBs (100 kb window). (D) Heatmaps showing the γH2AX (right panel) and BLM (left panel) ChIP-seq signal on a 100 kb window centered around DSB sites.

Individual inspection of DSBs revealed that for equivalent cleavage (determined by BLESS (Clouaire et al., 2018)), some DSBs displayed increased BLM binding compared to others (see an example Fig. 2A). Given the ability of BLM to bind G4, we further compared BLM binding with the available G4 mapping generated by BG4 immunoprecipitation in HaCaT cells (Hänsel-Hertsch et al., 2016). Interestingly following damage induction, BLM tended to be recruited at loci enriched in G4 (Fig. 2A). In order to determine the genomic/epigenomic determinants that foster BLM recruitment at some DSBs compared to others we further proceeded to identify subsets of DSBs differentially enriched in BLM, as previously described (Aymard et al., 2014; Clouaire et al., 2018). To this aim, we used our previously published BLESS dataset (Clouaire et al., 2018) to take into account unequal activity of AsiSI throughout the genome and computed the BLM/BLESS enrichment ratio for each of the 80 most cleaved DSBs (Material and Methods, (Clouaire et al., 2018)). This approach indeed allowed us to identify a subset of DSBs enriched in BLM (BLM-positive) while some others are only poorly able to recruit BLM (BLM-negative) (Fig. S1A-B). On average, BLM-positive DSBs were enriched in G4 compared to the BLM-negative subset of DSBs (Fig. 2B).

**Figure 2.**
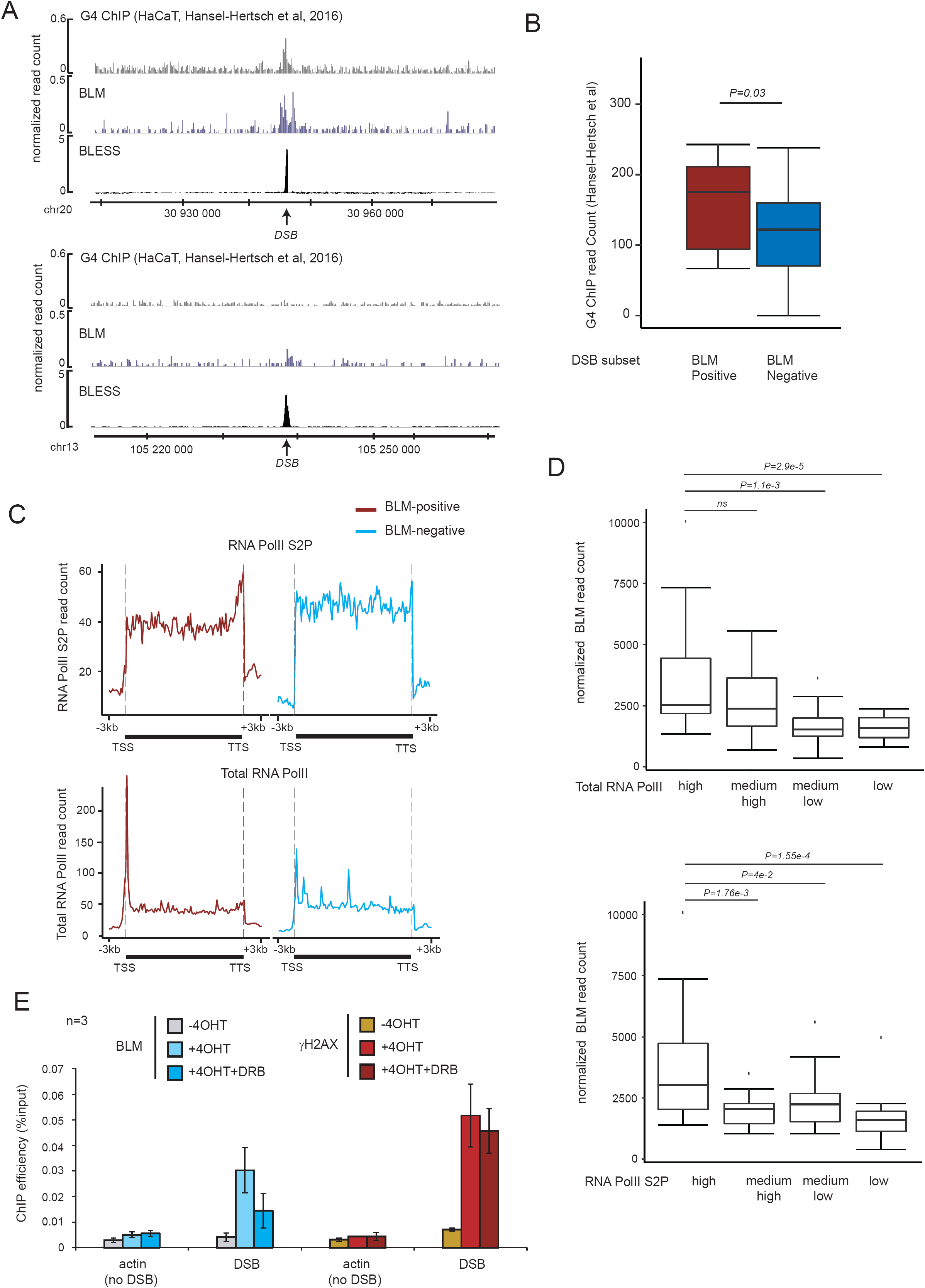
Transcription-dependent BLM recruitment at DSBs. (A) Genome browser screenshot representing BLM ChIP-seq, BLESS signal as well as G4 mapping (BG4 IP from (Hänsel-Hertsch et al., 2016)) at two DSBs cleaved at an equivalent level (chr20: 30946312, enriched in BLM and chr13:105238551, not enriched in BLM). (B) Boxplot representing G4 ChIP-seq read count (SRE5586987) (Hänsel-Hertsch et al., 2016) on a 1kb window for BLM-positive (red) and negative (blue) DSB sites. *P*=0.035 (Wilcoxon test). (C) Average ChIP-seq profiles of RNA Pol II-S2P (top panel) and Total RNA Pol II (bottom panel) across the closest genes lying near BLM-positive (red) or BLM-negative DSBs (blue). (D) Boxplot representing normalized BLM ChIP-seq read count after DSB induction on loci with high, medium high, medium low, or low RNA Pol II occupancy (upper panel) or RNA Pol II-S2P occupancy (lower panel) before DSB induction (determined by ChIP-seq before DSB induction). *P* values (Wilcoxon t-test) are indicated. (E) BLM (blue) and γH2AX (red) ChIP efficiency (expressed as % of input immunoprecipitated) before (−4OHT), after (+4OHT) DSB induction and after DSB induction with prior inhibition (1h pre-DSB induction) of RNA Pol II transcription elongation (+4OHT+DRB) at 80 bp from a DSB enriched in BLM (chr22:38864101) and on a control site (actin; no-DSB). Mean and SEM from three independent biological replicates are shown.

We further analyzed BLM binding with respect to transcriptional activity. Individually, DSB recruiting BLM tended to be in transcribed loci, compared to DSB displaying limited recruitment of BLM (Fig. S1C). We further used ChIP-Seq dataset of RNA-Polymerase II (total and phosphorylated on serine 2) generated before break induction in the DIvA model (Aymard et al., 2014; Cohen et al., 2018). The genes located at the immediate vicinity of BLM-positive DSBs exhibited the typical pattern of actively transcribed genes, with an accumulation of Total RNA pol II at promoters and RNA Pol II-S2P at transcription termination site (TTS) in contrast to the genes lying near BLM-negative DSBs (Fig. 2C). In agreement, loci that display strong RNA Pol II and RNA Pol II-S2P signal before DSB induction, displayed higher BLM recruitment following DSB induction (Fig. 2D). Altogether, these analyses revealed that BLM recruitment following DSBs is enhanced at loci showing G4 structures and transcriptional activity. To determine the influence of transcriptional activity on BLM recruitment, we next performed BLM ChIP in the presence of 5,6-Dichlorobenzimidazole 1-β-D-ribofuranoside (DRB), a selective inhibitor of RNA Pol II transcription elongation. BLM recruitment post damage (+4OHT) was strongly reduced in DRB pre-treated cells compared to untreated cells, while γH2AX signal, and hence DSB induction, was left unaffected (Fig. 2E), indicating that transcription activity fosters BLM recruitment upon DSB.

Hence altogether these data indicate that the ability of a DSB to recruit BLM depends on its genomic location, and more specifically on the transcriptional status of the underlying locus.

### BLM fosters resection, RAD51 binding and repair fidelity at DSB induced in transcriptionally active loci

Given that BLM has been extensively involved in HR, we next compared the binding of BLM with that of RAD51, previously reported by ChIP-seq in DIvA cells (Aymard et al., 2014). Notably, BLM strongly paralleled RAD51 distribution (see an example Fig. 3A and aggregate profiles at BLM-positive and BLM-negative DSBs subsets Fig. 3B). Moreover, BLM-negative sites displayed a low ability to recruit RAD51 (Fig. 3A-B), despite equivalent cleavage (BLESS signal, Fig. 3A). Of interest, mapping of RAD51 and BLM after longer exposure to AsiSI activity (24h) showed considerably extended spreading of RAD51 and BLM (Fig. 3C), with RAD51 distribution again highly correlating with BLM distribution (Fig. 3D). Altogether these data suggest that BLM spreads on surrounding chromatin as resection progresses and as the RAD51 filament assembles.

**Figure 3.**
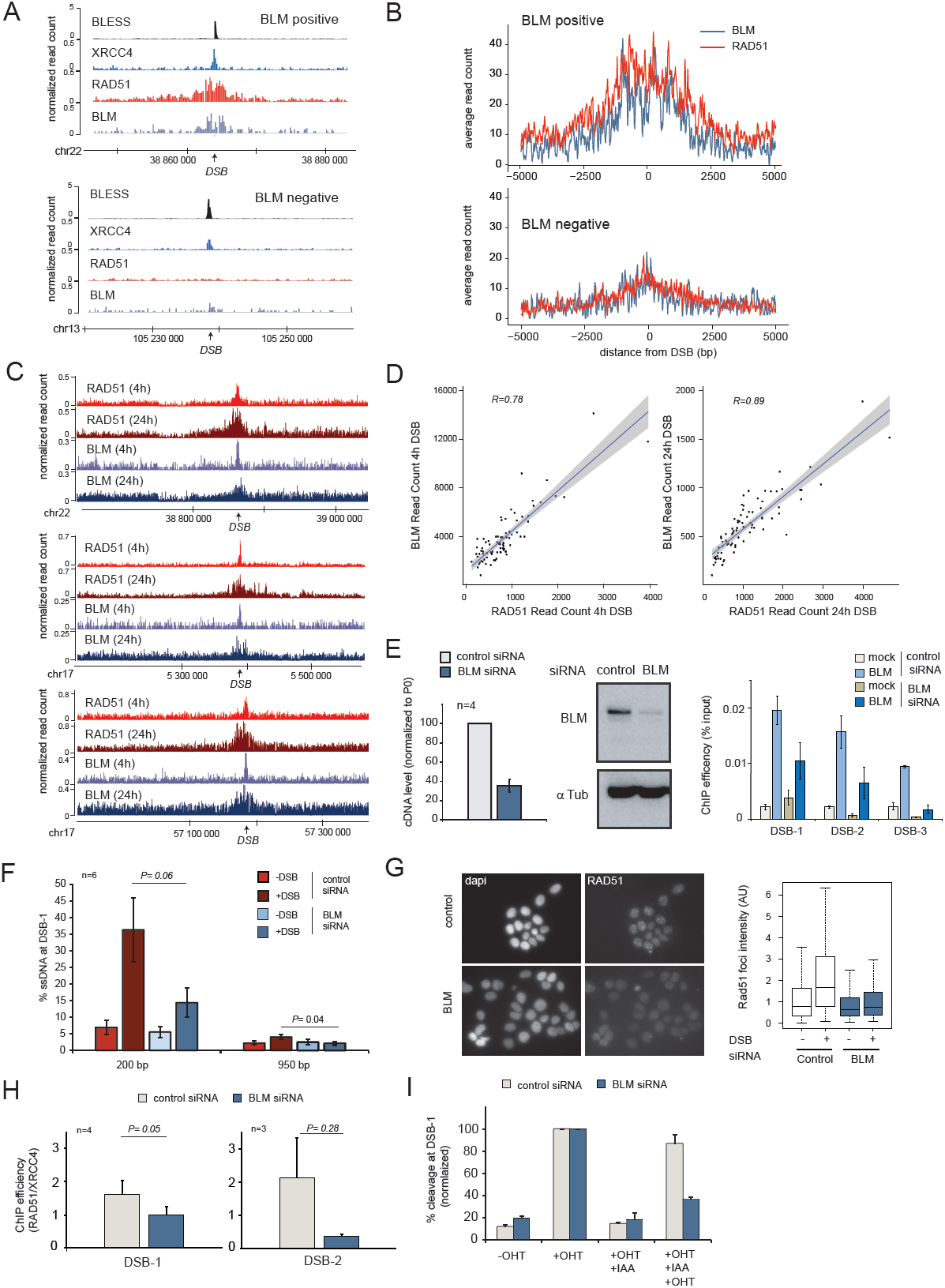
BLM fosters resection, RAD51 loading and repair fidelity. (A) Genome browser screenshot representing BLESS, XRCC4, RAD51 and BLM ChIP-seq signal at a BLM-positive DSB (upper panel; chr22:38864101) and a BLM-negative DSB (lower panel; chr13:105238551). DSBs are indicated by arrows. (B) Average profiles of BLM (blue) and RAD51 (red) on BLM-positive and BLM-negative DSB sites after DSB induction (10kb window). (C) Genome browser screenshot representing RAD51 and BLM ChIP-seq signal at 4hr and 24hr after DSB induction at three DSBs (arrows) (located on chr22:38864101; chr17:5390220 and chr17:57184296). (D) Scatter plot representing BLM and RAD51 read counts at the best eighty DSBs on a 10kb window at 4h (left panel) and on a 40kb at 24h (right panel) after DSB induction. (E) Left panel, normalized cDNA level in control and BLM siRNA transfected DIvA cells. Middle panel, immunoblot (using Tubulin and BLM antibodies) following BLM and control siRNA transfection as indicated. Right panel, BLM and mock (no antibody) ChIP in 4OHT-treated DIvA cells transfected with control and BLM siRNA as indicated. BLM enrichment was assessed by qPCR at three DSB showing BLM signal by ChIP-seq (DSB-1 chr22:38864101; DSB-2 chr9:130693170, DSB-3 chr20: 30946312) (F) Resection assay showing the percentage of single strand DNA (%ssDNA) in control (red) and BLM (blue) siRNA depleted DIvA cells at two distances from the DSB-1 (chr22:38864101, bound by BLM) as indicated. (G) (Left) RAD51 immunostaining performed in control and BLM siRNA depleted DIvA cells after DSB induction. (Right) Quantification of RAD51 foci intensity before (−) and after (+) DSB induction (> 100 nuclei). (H) ChIP efficiency (RAD51/XRCC4) at two DSBs (DSB-1 chr22:38864101 and DSB-3 chr20:30946312) in control (grey) and BLM (blue) siRNA depleted DIvA cells after DSB induction. (I) Cleavage efficiency showing the percentage of cleavage (normalized) in control (grey) and BLM (blue) siRNA depleted AID DIvA cells before DSB (−OHT), after DSB (+OHT), after repair (+OHT+IAA) and after a second round of DSB induction (+OHT+IAA+OHT) (DSB-1 on chr22:38864101).

In order to determine the function of BLM at these transcribed damaged loci, we performed siRNA depletion (Fig. 3E left panel and middle panel) and confirmed a clear drop in BLM recruitment on damaged chromatin (Fig. 3E right panel) without disturbing the cell cycle (Fig. S2A). Using a previously described assay that relies on the inability of restriction enzymes to cleave single-strand DNA (Zhou et al., 2014), we found that BLM depletion triggered a reduction in resection at a DSB bound by BLM (Fig. 3F), indicative of a pro-resection activity of BLM at DSB induced in active loci. In agreement, BLM depletion also triggered a strong reduction in RAD51 foci formation (Fig. 3G) and in RAD51 binding at DSBs detected by ChIP (Fig. 3H, Fig. S2B). Altogether this indicates that BLM is fostering resection and RAD51 binding at DSBs induced in active chromatin.

To assess repair fidelity in a BLM deficient background, we further used AID-DIvA cells. In this cell line, AsiSI-ER is fused to an auxin inducible degron (AID), allowing its rapid degradation upon auxin addition, thereby enabling the repair of AsiSI-induced DSBs (Aymard et al., 2014; Caron et al., 2015). Repair kinetics was globally left unaffected upon depletion of BLM, as measured by γH2AX foci resolution (Fig. S2C) or qPCR-based cleavage assay (Chailleux et al., 2014) (Fig. S2D), indicating that the lack of BLM, although reducing HR, does not delay DSB re-joining. In agreement, BLM depletion did not trigger any defect in cell survival following DSB induction, compared to control cells and if anything, it slightly increased cell recovery following DSB (Fig. S2E). Altogether these data indicate that BLM fosters HR usage at DSB induced in transcribed loci but its absence does not really impact the overall capacity of human cells to handle DSBs in active chromatin.

In this context, repair fidelity can be assessed. In case of accurate repair, the AsiSI restriction site is reconstituted post-auxin addition (OHT+IAA), and becomes thus available for a new round of cleavage by 4OHT treatment (Caron et al., 2015) (OHT+IAA+OHT). Interestingly, in BLM-depleted cells, AsiSI site cleavage efficiency was reduced after an additional round of 4OHT treatment compared to control cells (Fig. 3I) indicative of decreased repair fidelity upon BLM depletion.

Altogether these data indicate that at DSBs occurring in transcribed loci, BLM contributes to resection and RAD51 loading, to ensure faithful Transcription Associated Homologous Recombination Repair (TAHRR), the G2 arm of TC-DSBR, but that this function is not mandatory to ensure rapid end-rejoining and cell survival.

### BLM depletion rescues cell death in SETX-deficient cells following DSB in transcribed loci

We recently identified the SETX RNA:DNA helicase as a critical component involved in TC-DSBR, thanks to its ability to unwind RNA:DNA hybrids (Cohen et al., 2018; Puget et al., 2019). Alike BLM, SETX is specifically recruited at DSBs induced in transcribed loci and not at DSBs induced in intergenic and silent genes/promoters. Interestingly, we previously reported that SETX deficiency impairs RAD51 foci assembly and HR (Cohen et al., 2018). Given our above findings we hence further investigated the interplay between SETX and BLM in TC-DSBR. Notably, SETX and BLM spanned similar regions surrounding DSBs (Fig. 4A) and the recruitment level of both proteins significantly correlated at all DSBs (Fig. 4B). However unexpectedly, while as previously shown (Cohen et al., 2018) SETX impaired cell survival following DSBs induced in transcribing loci, BLM deficiency partially rescued the cell death observed in SETX-deficient cells (Fig. 4C). Yet, BLM knock-down was unable to rescue defective RAD51 foci formation in SETX depleted cells (Fig. 4D). Altogether this indicates that BLM depletion promotes cell survival in a SETX-deficient context where RNA:DNA hybrids accumulates, despite being unable to restore HR.

**Figure 4:**
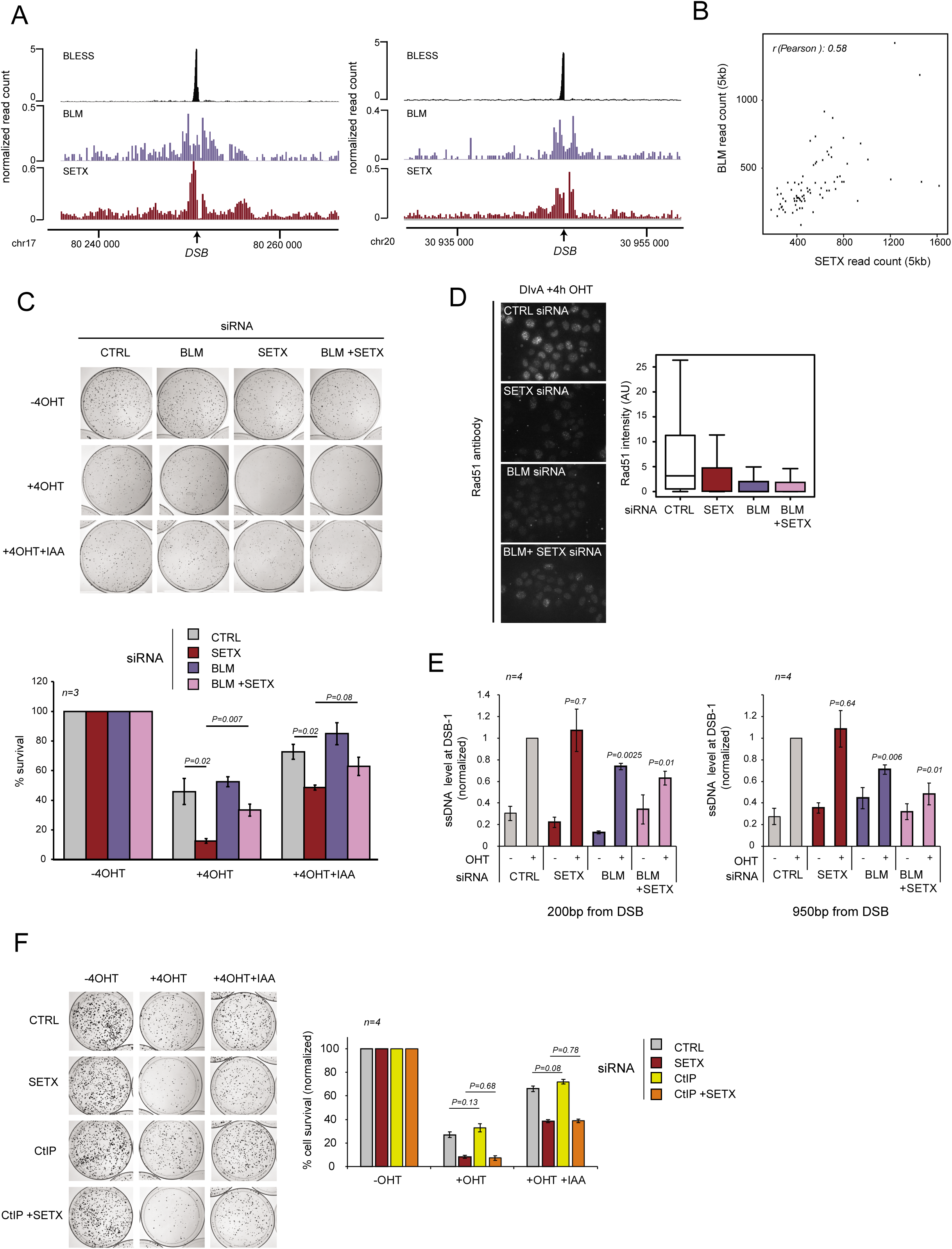
BLM deficiency rescues cell lethality in a context of accumulation of RNA:DNA hybrids at DSBs in transcribed loci. (A) Genome browser screenshot representing BLESS, BLM and SETX ChIP-seq signal at two individual DSBs (chr17:80250841 and chr20: 30946312) (arrows). (B) Scatter plot showing the ChIP-seq BLM and SETX read count read on a 5kb window around the eighty-best induced DSBs at 4h after DSB induction. (C) Clonogenic assay in control, BLM, SETX and BLM/SETX siRNA depleted AID DIvA cells before (−4OHT), after (+4OHT) DSB induction and after DSB induction and auxin (IAA) treatment allowing DSB repair (+4OHT+IAA) as indicated (upper panel). The quantification is represented by the percentage of cell survival in the lower panel. N=3 and P values are indicated (paired t-test). (D) (Left) RAD51 immunostaining performed in control, BLM, SETX, BLM/SETX siRNA depleted DIvA cells after DSB induction (DIvA+4h OHT). (Right) RAD51 foci intensity was quantified after DSB induction (> 100 nuclei). (E) Resection assay showing the level of single strand DNA (normalized on the control +DSB) in control, SETX, BLM, and BLM/SETX siRNA depleted DIvA cells at 200 bp and 950 bp from a DSB bound by BLM (DSB-1 chr22:38864101). *P*-values (one sample t-test, assessing significance with siRNA Ctrl +OHT ssDNA level are indicated) (F) Clonogenic assay in control, SETX, CtIP, and CtIP/SETX siRNA depleted AID DIvA cells before (−4OHT), after (+4OHT) DSB induction and after DSB induction and auxin (IAA) treatment allowing DSB repair (+4OHT+IAA) as indicated (left panel). Quantification (right panel) shows the percentage of cell survival (mean and SEM across N=4 experiments). *P* values are indicated (paired t-test).

We previously reported that, while impairing HR, SETX deficiency did not decreased ssDNA production at sites of damage (Cohen et al., 2018), a feature also recently reported in yeast (Rawal et al., 2020). SETX was thus proposed to contribute to HR downstream of resection but upstream of RAD51 nucleofilament assembly by dissolving RNA:DNA hybrids (Cohen et al., 2018). The DSB-induced lethality observed in SETX-deficient background may hence arise from the inability of repair pathways to handle DSBs that are in a resected state and unable to assemble RAD51 filament. Given that BLM promotes resection, we considered that it could hence rescue the lethality of SETX deficiency by simply reducing ssDNA production. In agreement with this hypothesis, resection was impaired in a double SETX/BLM-deficient background to an extent comparable to the single BLM depletion (Fig. 4E). However, CtIP depletion, which triggers strong resection deficiency at DSBs including in DIvA cells (Zhou et al., 2014) (Fig. S3A-B), was unable to rescue cell survival upon DSB induction in SETX-deficient cells (Fig. 4F). This indicates that reduction of ssDNA generation by itself is not sufficient to counteract the adverse effect of hybrids accumulation at DSBs on cell survival.

### BLM promotes break-induced replication in a SETX-deficient background

In *S. cerevisae*, R-loop removal deficient, triple knock out strains for *rnaseH1, rnaseH2* and *Sen1* (SETX yeast ortholog) initiate break-induced replication (BIR) at sites of R-loop-induced damages, with strong consequences on viability (Amon and Koshland, 2016; Costantino and Koshland, 2018). In order to test whether BIR could account for defective cell survival following DSB induction in active genes upon SETX knock-down in human cells, we depleted POLD3 (Fig. S3A), the human pol32 yeast ortholog, which has been previously involved in mammalian BIR (Costantino et al., 2014; Dilley et al., 2016). Importantly, codepletion of POLD3 rescued cell survival following DSB induction when compared to SETX-only deficient cells (Fig. 5A). A similar result was obtained for the Pif1 helicase, which is also required for BIR in yeast (Fig. 5B; Fig. S3A) (Saini et al., 2013; Wilson et al., 2013) (Fig. 5B). Altogether, these results suggest that the impaired cell survival observed upon DSB induction in a background deficient for RNA-DNA hybrids dissolution is due to BIR.

**Figure 5.**
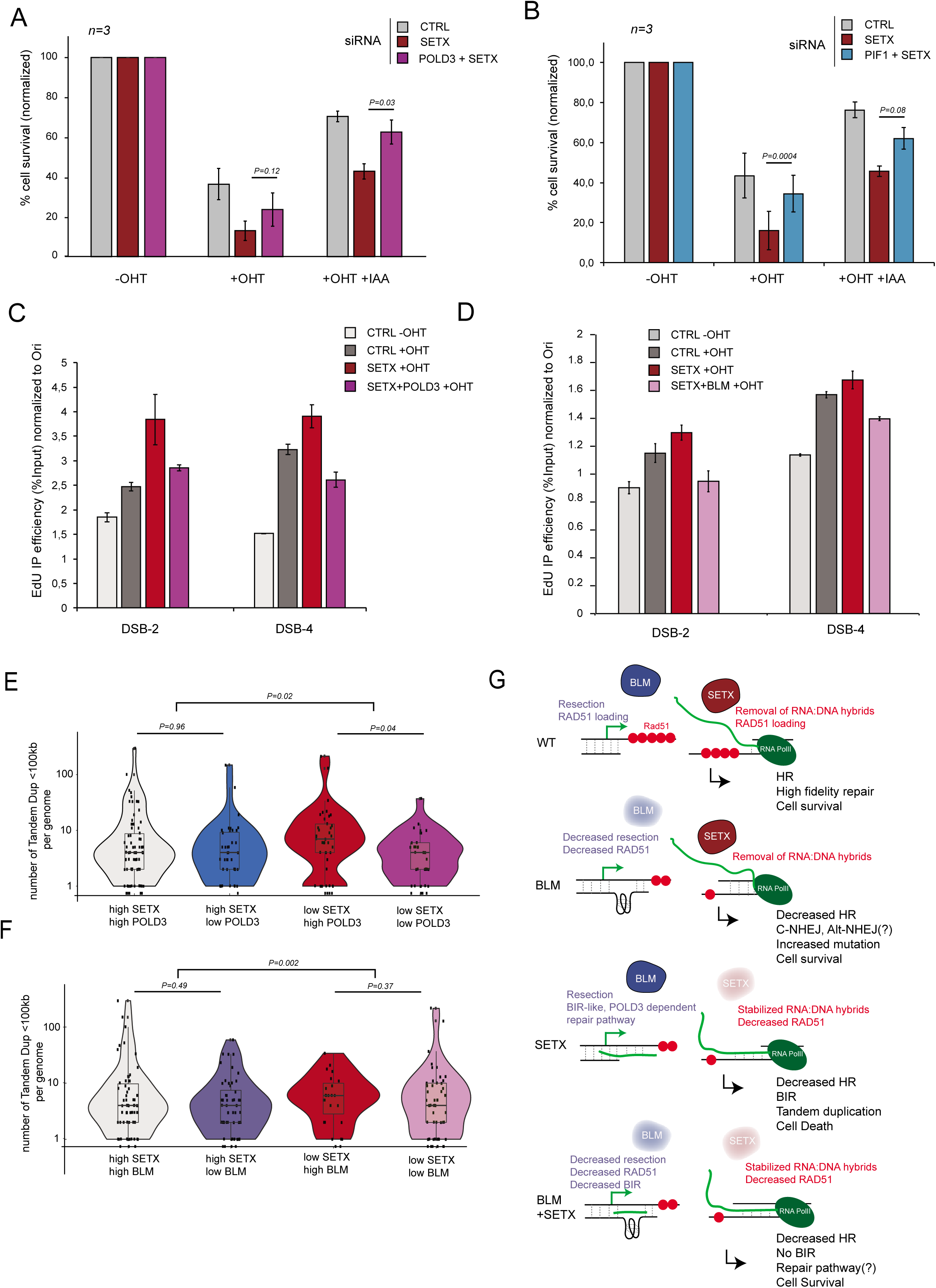
BLM promotes a BIR-like pathway in a SETX-deficient background. (A) Percentage of cell survival from clonogenic assay in control, SETX and POLD3/SETX siRNA depleted AID DIvA cells before (−4OHT), after (+4OHT) DSB induction and after repair (+4OHT+IAA). N=3. *P* values are indicated (paired t-test). (B) Percentage of cell survival from clonogenic assay in control, SETX and PIF1/SETX siRNA depleted AID DIvA cells before (−4OHT), after (+4OHT) DSB induction and after repair (+4OHT+IAA). N=3. *P* values are indicated (paired t-test). (C) EdU-IP efficiency in control, SETX and SETX/POLD3 siRNA depleted DIvA cells before and after DSB induction, at two DSBs bound by BLM (DSB-2, chr9:130693170, DSB-4, chr17:57184296). Data are normalized to an origin of replication (Ori). A representative experiment is shown. (D) EdU-IP efficiency in control, SETX and SETX/BLM siRNA depleted DIvA cells before and after DSB induction, at two DSBs bound by BLM (DSB-2, chr9:130693170, DSB-4, chr17: 57184296). Data are normalized to an origin of replication (Ori). A representative experiment is shown. (E) Violin Plots representing the number of tandem duplication (<100kb) per genome depending SETX and POLD3 expression as indicated using pancreatic cancer gene expression and genomic data available on ICGC database (PACA-CA project). *P* values between two groups (High SETX/High POLD3 with High SETX/Low POLD3, and Low SETX/High POLD3 with Low SETX/ Low POLD3) are indicated (unpaired Wilcoxon’s test). *P* value assessing the significance between ratios (whether “high SETX” group behaves differently than “Low SETX” group) is also indicated. (F) Violin Plots representing the number of tandem duplication (<100kb) per genome depending SETX and BLM expression as indicated using pancreatic cancer gene expression and genomic data available on ICGC database (PACA-CA project). *P* values between two groups (High SETX/High BLM with High SETX/Low BLM, and Low SETX/High BLM with Low SETX/ Low BLM) are indicated Wilcoxon’s test (unpaired). *P* value assessing the significance between ratios (whether “high SETX” group behaves differently than “Low SETX” group) is also indicated. (G) Schematic representation of the function of SETX and BLM in TC-DSBR. In WT conditions, SETX removes the RNA:DNA hybrids that accumulate at DSBs induced in transcribed loci through a yet unclear mechanism. BLM-dependent end-resection and RAD51 loading occur to promote faithful DSB repair by HR and cell viability. In BLM-deficient cells, end-resection and RAD51 loading are impaired which favors mutational repair pathways while maintaining cell viability. In SETX depleted cells, end resection occurs but deficient RNA:DNA hybrids removal reduces RAD51 loading and HR repair. DSB are further handled by a BLM, POLD3 BIR-like pathway, triggering an increase of tandem duplications on the genome and severely impairing cell survival. In BLM/SETX-depleted cells, RNA:DNA hybrids are stabilized, impeding RAD51 loading and HR, but the absence of BLM impairs BIR and as a consequence increases cell viability. The DSB repair pathway used in these cells remains unknown (C-NHEJ, Alt-NHEJ?).

Since BIR implies new DNA synthesis, we further set out to measure DNA synthesis at sites of DSBs. For this we implemented a method to directly purify newly synthetized DNA following DSB induction using *in vivo* EdU incorporation, followed by click chemistry and purification. In control cells, we could readily detect 4OHT-induced, repair-associated DNA synthesis at the vicinity of DSBs (Fig. S3C), compared to replicative synthesis at an origin of replication. In SETX-deficient cells where RAD51 filament assembly and hence HR-driven DNA synthesis are impaired, we nevertheless detected increased DNA synthesis at break sites (Fig. S3D, Fig. 5C). This DNA synthesis was dependent on POLD3, indicating that BIR indeed occurs in the context of inefficient removal of RNA:DNA hybrids at sites of damage (Fig. 5C). Importantly, similar to POLD3, BLM depletion also decreased DNA synthesis in SETX-deficient cells (Fig. 5D) suggesting that BLM is necessary for activation of BIR in that context. Altogether these data suggest that upon excessive accumulation (impaired removal) of RNA:DNA hybrids at DSBs in transcribed loci, BIR is initiated in a BLM-dependent manner and reduces cell survival.

BIR genomic signature entails copy number alteration, especially tandem duplication (TD) below 100kb (Costantino et al., 2014). We thus further investigated the status of TD<100kb in relationship with the expression levels of SETX. This was achieved using pancreatic cancer gene expression and genomic data available on ICGC database (PACA-CA dataset) since this project provides a robust set of patient samples in which TD have been annotated. Individual patient genomic data were categorized according to their SETX and POLD3 expression level (i.e. four categories as indicated, Fig. 5E). Interestingly, in condition where POLD3 expression is high, TD frequency was increased in samples showing low SETX expression compared to samples showing high SETX expression (Fig. 5E, compare red and grey). This indicates that BIR genomic signature increases in a R-loop dissolution deficient background. However, reduced expression of POLD3 significantly decreased TD in samples that display low expression of SETX (Fig. 5E, compared red and purple). Similarly, reduced BLM expression also decreased TD in patient samples showing low SETX expression (Fig. 5F, compare red and purple). Altogether this suggests that the BIR genomic signature that arise in pancreatic cancer cells displaying low R-loop unwinding abilities (low SETX expression) depends on POLD3 and BLM, further validating the previously observed genetic interaction between SETX, POLD3 and BLM and suggesting that the interplay between SETX, POLD3 and BLM directly impacts genome stability *in vivo*.

## Discussion

Here, we present evidence that the RecQ BLM helicase is a *bona fide* component of the TC-DSBR machinery and displays a dual role in handling DSBs that occur in actively transcribed loci. We unambiguously determined that BLM preferentially associates with DSBs located in active transcription units, where it fosters resection and RAD51 assembly. Yet in a background where R-loop dissolution is impaired, BLM displays a pro-BIR function, thereby decreasing cell viability (Fig. 5G).

### Function of BLM in TAHRR, the G2 arm of TC-DSBR

We found that BLM preferentially associates with DSBs localized in actively transcribed genes in a manner that depends on transcription (Fig. 2). This behaviour is reminiscent of other recently identified TC-DSBR factors, such as SETX (Cohen et al., 2018) or XPG and RAD52 (Yasuhara et al., 2018), whose recruitment at DSBs were also shown to depends on pre-existing transcriptional activity. In agreement with previous reported functions of BLM during HR, we found that at these DSBs in active loci, BLM contributes to resection, RAD51 foci assembly and accurate repair. Importantly, the observation that BLM spreads on surrounding chromatin as a matter of time and parallels RAD51 distribution suggests that BLM translocates along the DNA as resection progresses, subsequently favouring RAD51 loading. One of the well-established molecular activity of the BLM helicase is to unwind G4 (Chatterjee et al, 2014; Huber et al, 2002; Mohaghegh et al, 2001) and Strand-seq experiments indicated that BLM function is particularly critical to counteract SCE at G4-enriched transcribed loci (van Wietmarschen et al., 2018). Of importance, BLM was also reported to unwind G4 at DSBs (Day et al., 2017) and at ROS-induced DNA damage (Tan et al., 2020). Furthermore, another G4 helicase, Pif1 was shown to promote resection thanks to its ability to remove G4 (Jimeno et al., 2018). Given that G4 are prevalent in transcribed regions and especially gene promoters (Chambers et al, 2015; Huppert & Balasubramanian, 2007), and that BLM binding displays a bias for G4-enriched DSBs (Fig. 2), a tempting hypothesis is that at these DSBs, BLM functions thanks to its G4-helicase activity to allow progression of resection (Fig. 5G).

### BLM, and hence TAHRR, is dispensable for DNA end re-joining and cell survival after DSB in active loci but favours accurate repair

Previous studies led to conflicting results regarding the sensitivity of BLM-deficient cells to DNA damaging agents. Indeed, both Bloom Syndrome (BS) and BLM-depleted cells were found to be insensitive to HU-induced replication stress, unless prolonged HU treatment is performed (Davies et al, 2004; Lahkim Bennani-Belhaj et al, 2010). Similarly, BLM-defective chicken DT40 cell exhibit normal sensitivity to HU (Imamura et al, 2001). In contrast, BS and/or BLM siRNA depleted cells display hypersensitivity to formaldehyde (inducing DNA-protein cross-links, (Kumari et al, 2015)), agents inducing DNA interstrand cross-links (Pichierri et al, 2004) camptothecin (Rao et al, 2005) or other genotoxics (for example see (Beamish et al, 2002; So et al, 2004)). Here, we unambiguously demonstrate that the transient depletion of BLM does not reduce survival of cells that experience a hundred of clean DSBs, a majority of them being induced in active loci.

In agreement with this, we found that BLM depletion did not impair the kinetics of DSB ends rejoining. This is consistent with several studies showing that end joining is not reduced in BS cells or nuclear extracts, at least when using plasmids as substrates (Gaymes et al, 2002; Langland et al, 2002; Onclercq-Delic et al, 2003). However, although not necessary for repair, its depletion decreased accuracy of the repair process. Our data would be in agreement with Alt-NHEJ usage, previously proposed to contribute to DSB repair in BLM-deficient cells (Adams et al, 2003; Gaymes et al, 2002; Grabarz et al, 2013; Wang et al, 2011). Interestingly, plasmids-based assays have failed to detect unfaithful repair in BS cells (Langland et al, 2002; Onclercq-Delic et al, 2003), further strengthening the idea that BLM may be required to promote repair accuracy in very specific chromatin contexts such as active genes.

More importantly, while HR is the preferred pathway that handles DSBs in transcribed chromatin during G2 (Aymard et al., 2014), our data indicate that HR usage is not required to ensure cell viability in this context. This is in agreement with the recent finding that CRISPR/Cas9 induced breaks in DNAse hypersensitive sites (open chromatin) are prone to resection and that inhibiting resection at these DSBs actually improves cell survival (van den Berg et al., 2019). Hence altogether this suggests that DSB in transcribed loci mainly utilizes HR (TAHRR) to ensure accurate repair, but that TAHRR deficiency is not cytotoxic while nevertheless affecting genome stability. Given our results that BLM mainly functions at damaged active genes, it is tempting to speculate that the cancer predisposition observed in BS patients arise at least in part from inaccurate repair events at damaged active transcription units, which would hence undergo mutagenic repair while still being proficient for proliferation.

### TC-DSBR entails a BIR-like mechanism when RNA:DNA hybrids resolution is impaired

Previous work indicated that, once damaged, transcribed loci accumulate RNA:DNA hybrids and that their dissolution is mandatory for the proper execution of HR (Cohen et al., 2018; Yasuhara et al., 2018). Here we report that in mammalian cells, a BLM-dependent BIR-like pathway handles such DSBs in actively transcribed loci when R-loop removal is impaired, in agreement with previous studies in yeast showing that DSBs are repaired by BIR in strains accumulating R-loops (Δ *rnaseh1, rnaseh2* and *sen1*) (Amon and Koshland, 2016; Costantino and Koshland, 2018). Indeed we found that (i) the depletion of POLD3 rescues lethality upon DSB in active genes in a SETX-depleted background, (ii) excessive DNA synthesis takes place at DSBs upon SETX depletion despite reduced RAD51 loading (Cohen et al., 2018), (iii) this repair-associated DNA synthesis depends on POLD3 and BLM, and (iv) BIR genomic signature correlates with expression levels of SETX, POLD3 and BLM in cancer patient samples. So far, POLD3-dependent BIR in human cells has been mainly involved in Alternative Lengthening of Telomeres (ALT) (Dilley et al., 2016; Min et al., 2019; Porreca et al., 2020; Roumelioti et al., 2016; Sobinoff et al., 2017; Zhang et al., 2019) and at one-ended DSBs, collapsed replication forks by a process also known as MiDAS (Mitotic Induced DNA synthesis) (Bhowmick et al., 2016; Costantino et al., 2014; Minocherhomji et al., 2015; Sotiriou et al., 2016). Moreover, BIR has been previously proposed to account for specifics genomic signatures, including tandem duplication, on the human genomes (Carvalho et al., 2013; Costantino et al., 2014; Hastings et al., 2009a, 2009b; Willis et al., 2015; Zhang et al., 2009), suggesting that BIR indeed occurs in some context in mammalian cells.

Here, we provide evidence that such a BIR-like pathway can operate at intra-chromosomal two ended-DSBs. Though granting further studies, we speculate that the R-loop accumulation that takes place at DSBs induced in active loci may actually convert a two-ended DSB into a “one-ended-like” DSB (through for instance asymmetrical RNA:DNA hybrids accumulation).

Importantly alike to what is described here, ALT-BIR pathway not only depends on BLM (Sobinoff et al., 2017; Zhang et al., 2019) but also increases upon impairment of R-loops removal (Lu et al., 2019; Pan et al., 2017, 2019; Silva et al., 2019). In a similar manner, BIR requires Sgs1 (the BLM ortholog) in yeast and is fostered by Mph1, the FANCM yeast ortholog (Mehta et al., 2017), an helicase known to display RNA:DNA hybrids unwinding activity (Schwab et al., 2015). Altogether this suggests that R-loop accumulation is central to the BIR process, both in yeast and mammalian cells. In addition during ALT, telomere clustering fosters BIR (Loe et al., 2020; Min et al., 2019). Interestingly, we previously reported that DSBs induced in active loci display clustering (Aymard et al., 2017). Whether the BIR pathway described here that handles DSB in active genes upon impairment of R-loop removal also relies on their ability to cluster deserves further investigations.

Altogether our work indicates that during DSB repair, RNA:DNA hybrids are highly toxic intermediates that potentiate the use of the cytotoxic BIR pathway. In this context, chemical compounds stabilizing R-loops in cancer backgrounds where the BIR pathway is intact may reveal a good strategy to enhance the potency of gene-specific genotoxic chemotherapeutic drugs (such a Topo II poison).

## Material and Methods

### Cell culture

DIvA (AsiSI-ER-U20S) and AID-DIvA (AID-AsiSI-ER-U20S) cells were cultured in Dulbecco’s modified Eagle’s medium (DMEM) supplemented with antibiotics, 10% FCS (InVitrogen) and either 1 µg/mL puromycin (DIvA cells) or 800 µg/mL G418 (AID-DIvA cells) at 37°C under a humidified atmosphere with 5% CO2. For AsiSI-dependent DSB induction, cells were treated with 300 nM 4OHT (Sigma; H7904) for 4 h. When indicated, 4OHT treated cells were washed 3 times with pre-warmed PBS and further incubated with 500 µg/mL auxin (Sigma; I5148). For transcriptional inhibition, DRB (100 μM) was added to the medium 1 h prior to 4OHT (4 h) treatment. siRNA transfection was performed using the *Amaxa®* Cell Line *Nucleofector® Kit V*, and program X001for 48h hours followed by a 4h 4OHT treatment. siRNA sequences are detailed in the Supplemental Table S1.

### Clonogenic assays

After siRNA transfection, AID-DIvA cells were seeded at a clonal density in 10 cm diameter dishes. 48h later cells were treated with 300 nM 4OHT for 4 h and, when indicated, washed 3 times with pre-warmed PBS and further incubated with 500 µg/mL auxin for another 4 h. After 3 washes with pre-warmed PBS, complete medium was added to each dish. After 7 to 10 days, cells were stained with crystal violet (Sigma) and counted. Only colonies containing more than 50 cells were scored.

### Chromatin immunoprecipitation followed by qPCR and high throughput sequencing

ChIP assays were carried out according to the protocol described in (Iacovoni et al., 2010) with the following modifications. 200 µg of chromatin was immunoprecipitated by using 2 µg of anti-BLM (Abcam, ab2179), anti-XRCC4 (Abcam, ab145), anti-RAD51 (Santa Cruz, SC-8349) or without antibody (mock). For quantitative PCR analysis, both input and IP samples were analyzed using the primers described in Supplemental Table 2. IP efficiencies were calculated as the percent of input DNA immunoprecipitated.

### Sequencing data analyses

For BLM and RAD51 ChIP-Seq after 4h and 24h 4OHT treatment, sequencing libraries were prepared by using 10 ng of purified DNA (averaged size 250-300 bp) and subjected to high throughput sequencing (single-end) using a HiSeq 2000 sequencing (BGI, Hong-Kong) or using Illumina NextSeq 500 (EMBL Genomics core facilities Heidelberg, Germany). ChIP-Seq experiments were aligned using bwa on human reference genome hg19. Samtools was used for sorting and indexing, and a custom R script with R package rtracklayer was used to generate normalized (CPM) coverage of mapped reads in bigWig format. ChIP-Seq average profiles were computed using normalized number of reads around the 80 best cleaved DSBs (Clouaire et al., 2018). For heat maps representation (performed usin Java Treeview (http://www.jtreeview.sourceforge.net) each tile shows the average normalized number of reads at each genomic location centered around DSBs, ordered based on the BLESS signal (Fig. 1D). For boxplots, the sum of the normalized number of reads of ChIP-Seq was computed on a given window around each DSB (as indicated). Box-plots were generated with R-base. The center line represents the median, box ends represent respectively the first and third quartiles, and whiskers represent the minimum and maximum values without outliers. Outliers were defined as first quartile − (1.5 × interquartile range) and above third quartile + (1.5 × interquartile range). Statistical hypothesis testing was performed using nonparametric paired Mann-Whitney-Wilcoxon (wilcoxon.test() function in R) to tests distribution differences between two populations. For comparison with RNA Pol II ChIP-seq data (Fig. 2D), each RNA Pol II categories were computed using coverage of normalized reads counts (ChIP-seq data in DIvA cells from (Cohen et al., 2018)) from a given windows of 10kb around each DSBs and divided in four groups using quartiles.

To determine categories of DSBs enriched in BLM or not, for each cleaved DSBs, BLESS (Clouaire et al., 2018) and BLM normalized coverage was computed using similar method used for boxplots, using a 1kb and 4kb window respectively. Then the ratio between these two values was computed and used to sort DSBs. The first 20 DSBs was considered as BLM-positive and the last 20 as BLM-negative. For comparison with G-quadruplexes (Fig. 2B), BG4-Seq files were downloaded from GSE99205 (Hänsel-Hertsch et al., 2016) using SRA6toolkit and fastq-dump command line, then processed using a classical ChIP-seq pipeline: bwa for mapping, samtools for sorting and indexing, and R custom script to generate normalized (CPM) coverage of mapped reads in bigWig format.

RNA-Seq and ChIP-Seq data for SETX, RNA Pol II total, RNA pol II phosphoS2 CTD were published previously (Cohen et al., 2018) and are available at ArrayExpress (E-MTAB-6318). Data for γH2AX ChIP-seq (Aymard et al., 2014) are available at ArrayExpress (E-MTAB-1241). BLESS data (Clouaire et al., 2018) are available at ArrayExpress (E-MTAB-5817). BLM and RAD51 ChIP-seq data generated in this study were deposited on Array Express (E-MTAB XXXX)

### Repair kinetics and repair fidelity at AsiSI sites

Repair kinetics at specific AsiSI-induced DSBs were measured as described in (Aymard et al, 2014) by a cleavage assay permitting the capture of unrepaired DSBs, at the indicated times after auxin addition. For fidelity assays, AID-DIvA cells were treated with 4OHT to induce DSBs for 4h (300nM) followed by an auxin treatment for 4h. The next day, cells were treated again with 4OHT for 4 h. DNA was extracted and subjected to a cleavage assay as described above. Primers used are detailed in the Supplemental Table 2.

### Immunofluorescence

Immunofluorescence against γH2AX (JBW301), and RAD51 (Santa Cruz) in AID-DIvA cells was performed as in (Cohen et al., 2018). Image acquisition was performed using MetaMorph on a wide-field microscope equipped with a with a cooled charge-coupled device camera (CoolSNAP HQ2), using a X40 objective (for quantification). Quantification was performed using Columbus, an integrated software to the Operetta automated high-content screening microscope (PerkinElmer). Dapi stained nuclei were selected according to the B method and appropriate parameters, such as the size and intensity of fluorescent objects, were applied to eliminate false-positive. Then γ-H2AX or RAD51 foci were detected with the D method with the following parameters: detection sensitivity, 1; splitting coefficient, 1; background correction, >0.5 to 0.9.

### Resection Assay

Measure of resection was performed as described in (Zhou et al, 2014) with the following modifications. DNA was extracted from fresh cells using the DNAeasy kit (Qiagen) and directly digested using the Ban I restriction enzyme that cuts at approximately 200bp and 950bp from a DSB located on chromosome 22 and enriched in BLM (chr22:38864101). Digested and undigested DNA were analysed by qPCR using the primers described in the Supplemental Table 2.

### Site specific DNA synthesis quantification by EdU-Immunopreciptation

After siRNA transfection by electroporation 4·10^6^ cells were seeded in 15 cm diameter dishes. After 48hr, cells are treated with 300 nM 4OHT (Sigma; H7904) for 4h and EdU (5-ethynyl-2’-deoxyuridine from Invitrogen; 10µM final) was added 1h before cells were harvested. DNA was extracted and digested with MboI (125U per 100µl reaction volume) for 12h at 37°C followed by NlaIII (12.5U per 100µl reaction volume) for 4h at 37°C. Digested DNA was purified with Phenol/chloroform (Sigma) and precipitated with ethanol (Sigma) and sodium acetate (Gibco). DNA was then biotinylated by click-it reaction according to the manufacturer protocol (Click-iT® Nascent RNA Capture Kit; Life technology) and precipitated with EtOH (Sigma) and Na-acetate (Gibco). Input DNA was taken at this step. Biotinylated DNA was immuno-precipitated using streptavidin beads according to manufacturer protocol (Click-iT® Nascent RNA Capture Kit; Life technology). Immuno-precipitated and input DNA were analyzed by qPCR using the primers indicated in the Supplemental Table 2.

### ICGC cancer data analysis

Publicly available cancer data from ICGC consortium data portal release 28 was retrieved (https://dcc.icgc.org/), with a focus on PACA-CA cohort (pancreatic cancer) for which tandem duplication mutations were available in enough patients for statistical analyses. For each patient of the cohort, the number of somatic tandem duplications whose sizes were shorter than 100 kb (from structural somatic mutation data) was computed. The patients were next divided into 4 groups depending on the expression of 2 genes SETX and polD3 or BLM: patients with high SETX expression and high PolD3 or BLM expression, patients with high gene SETX expression and low gene polD3 or BLM expression, patients with low gene SETX expression and high gene polD3 or BLM expression, and patients with low gene SETX expression and low gene polD3 or BLM expression. The number of tandem duplications between two groups was compared by a Wilcoxon’s test (unpaired). To compute the effect of one gene on the number of tandem duplications depending on the other gene, we used a negative binomial regression with an interaction term between the two genes and computed the corresponding p-value.

## Acknowledgments

This work was initiated by a CEFIPRA collaborative grant (4603-1). Funding in GL laboratory is provided by grants from the European Research Council (ERC-2014-CoG 647344), Agence Nationale pour la Recherche (ANR-14-CE10-0002-01) and the Ligue Nationale contre le Cancer (LNCC). S.C was supported by a fellowship from the Fondation pour la recherche medicale (FRM). VB laboratory was funded by grants from Agence Nationale pour la Recherche (ANR-15-CE11-0011 and ANR-18-CE12-0018), from Electricité de France, and from Fondation pour la Recherche Médicale (Equipe FRM EQU201903007785). T.C and N.P are INSERM researchers.

## Authors contributions

S.C., A.G., T.C., A.M., C.A, and N.P. performed and analyzed experiments. M.A., V.R., R.M. and M.G. performed bioinformatic analyses of ChIP-seq data sets and ICGC database. V.D. optimized EdU IP under the supervision of V.B. A.M., T.C., N.P., and G.L. conceived and supervised experiments. G.L. wrote the manuscript. All authors commented and edited the manuscript.

## Competing Interest

The authors declare no competing interest

## Notes

### Competing Interest Statement

The authors have declared no competing interest.

