## Supplementary material for "BLM-dependent Break-Induced Replication handles DSBs in transcribed chromatin upon impaired RNA:DNA hybrids dissolution": Supplemenraty File

Supplementary Figures legends

Figure S1-S3

Supplemental Table 1-2

**Figure S1.**

(A) Boxplot showing the ratio BLM/BLESS signal on BLM-positive (left) and BLM-negative (right) DSB sites in treated DIvA cells.

(B) Average profiles of BLM on BLM positive (red) and negative (blue) DSB (10kb window).

(C) Genome Browser screenshot of the BLM ChIP-seq signal (blue) in 4OHT-treated DIvA cells (post DSB induction), together with the RNA-seq (light green) and total RNA PolII ChIP-seq (green) signals obtained in DIvA cells prior DSB induction (Cohen et al., 2018) at five DSBs. DSBs 1-4 are further used in this study.

**Figure S2.**

(A) Cell cycle distribution of control and BLM siRNA-depleted DIvA cells as indicated. Mean and SEM are indicated for *N=3*.

(B) RAD51 (left panel) and XRCC4 (right panel) ChIP efficiency following treatment with siRNA control and BLM as indicated, at a DSB identified as bound by BLM (DSB-1, chr22:38862102). Mock ChIP was performed without antibody. A representative experiment is shown.

(C) Left panel, γH2AX immunostaining performed in control (control siRNA) and BLM (BLM siRNA)-depleted AID-DIvA cells after DSB induction (+4h OHT), after 30 min (+4h OHT+IAA 30') and 120 min (+4h OHT+IAA 120') of repair. Right panel, γH2AX foci intensity was quantified in the indicated conditions (> 100 nuclei).

(D) Cleavage efficiency showing the percentage of cleavage in control (grey) and BLM (blue) siRNA depleted AID DIvA cells before DSB (-4OHT), after DSB (+4OHT) induction, after 15min (+4OHT+IAA15') and 60 min (+4OHT+IAA60') of repair, as indicated, at a DSB identified as bound by BLM (DSB-1, chr22:38862102).

(E) Clonogenic assay in control and BLM siRNA depleted AID DIvA cells before (-4OHT), after (+4OHT) DSB induction and after repair (+4OHT+IAA) as indicated (upper panel). Quantification (lower panel) is represented by the percentage of cell survival. Mean and SEM across N=3 experiments are shown. *P* values are indicated (paired t-test).

**Figure S3.**

(A) Normalized cDNA level showing CtIP (left panel), PIF1 (middle panel) and POLD3 (right panel) siRNA-mediated knockdown efficiency

(B) Resection assay showing the level of single-strand DNA (normalized on the control +DSB) in control and CtIP-depleted DIvA cells at 200 bp from DSB-1, bound by BLM (chr22:38864101). A representative experiment is shown.

(C) EdU-IP efficiency before (-DSB) and after (+DSB) DSB induction on an origin of replication (Ori; used as a control region to measure the background of DNA synthesis due to DNA replication) at two DSBs bound by BLM (DSB-2, chr9:130693170, DSB-4, chr17: 57184296). A representative experiment is shown.

(D) Repair synthesis represented by the ratio +/- DSB of EdU-IP efficiency on a -/+ 3kb region around the DBS-2 (chr9:130693170) (normalized on the control region (Ori)) in control and SETX siRNA-depleted DIvA cells as indicated. Mean and SEM of N=3 are shown.

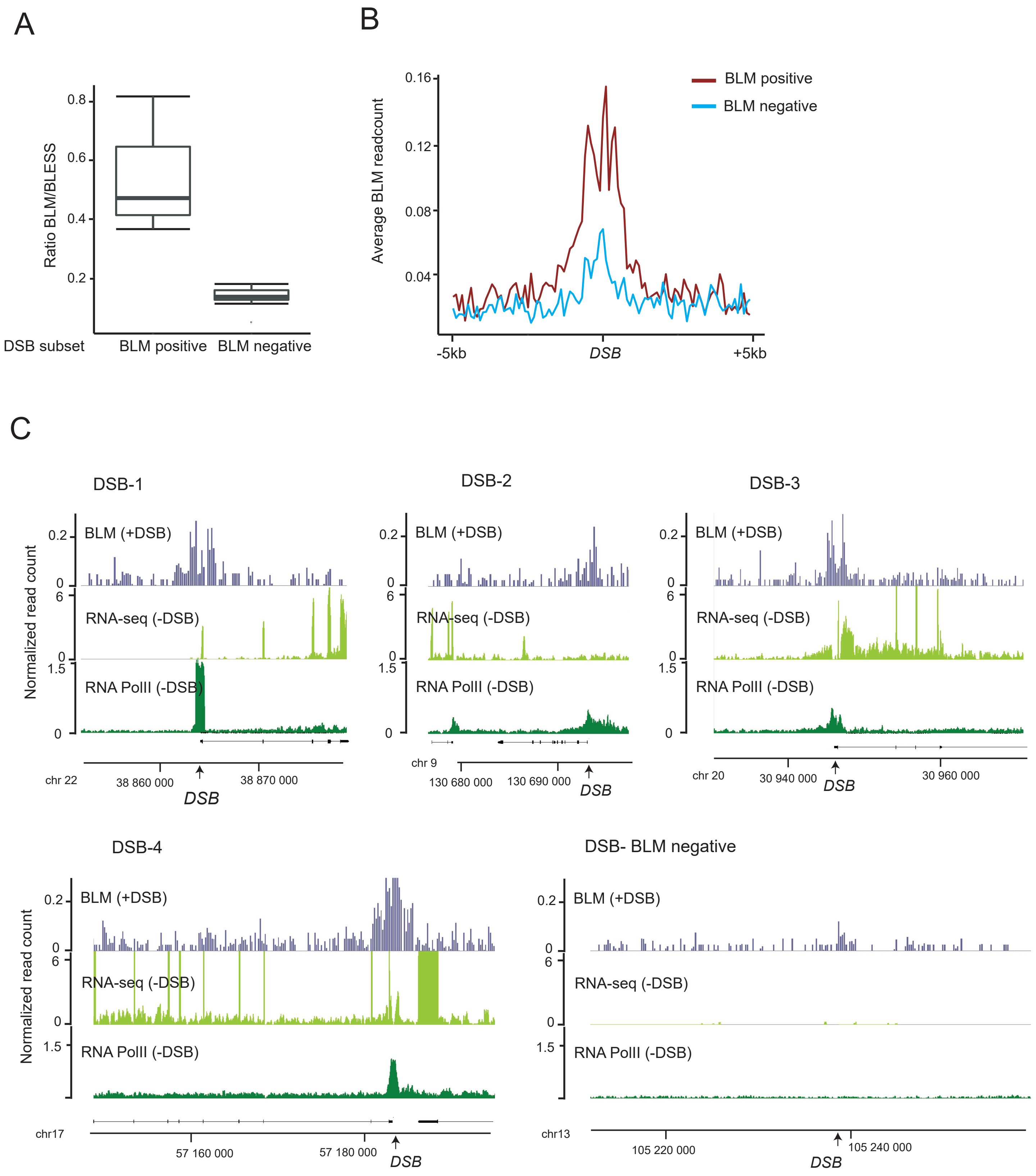
A
Ratio BLM/BLESS
0.8
0.6
0.4
0.2
DSB subset
BLM positive
BLM negative
B
Average BLM readcount
0.16
0.12
0.08
0.04
BLM positive
BLM negative
-5kb
DSB
+5kb
C
DSB-1
DSB-2
DSB-3
Normalized read count
BLM (+DSB)
RNA-seq (-DSB)
RNA PolII (-DSB)
chr 22
38 860 000
38 870 000
DSB
chr 9
130 680 000
130 690 000
chr 20
30 940 000
30 960 000
DSB-4
DSB- BLM negative
chr17
57 160 000
57 180 000
chr13
105 220 000
105 240 000

Cohen, Guenolé et al, Figure S2

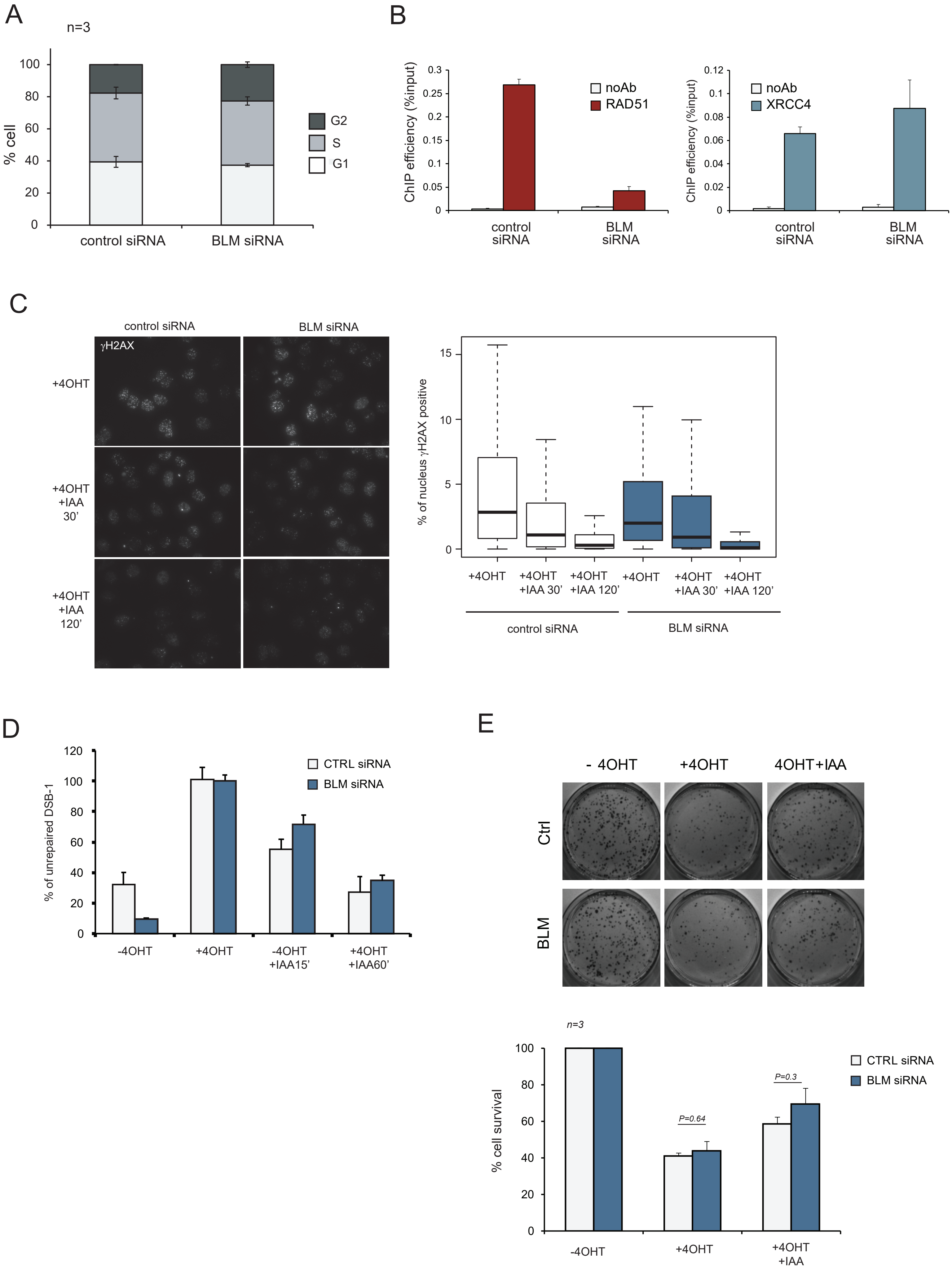

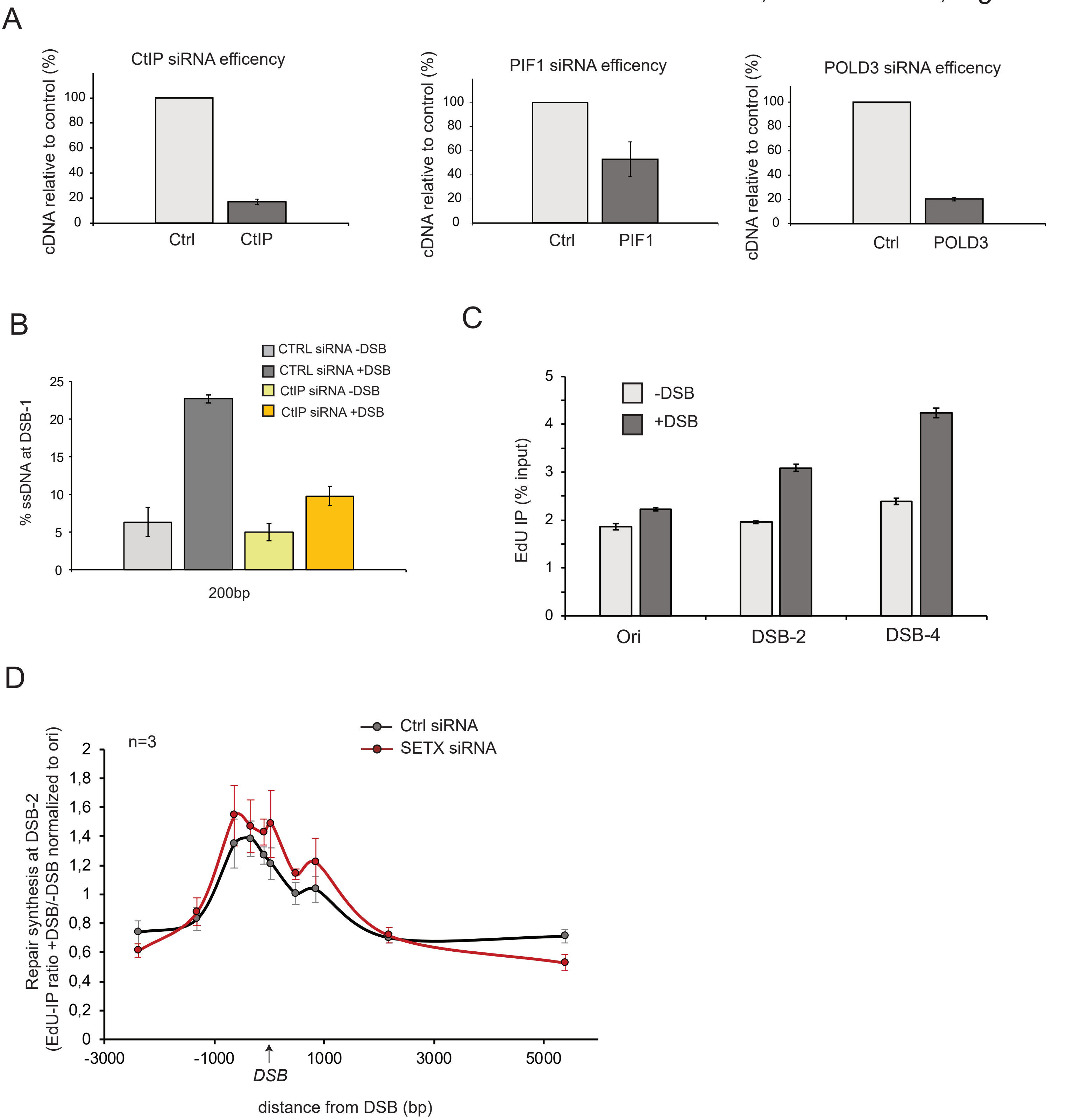
A
CtIP siRNA efficency
cDNA relative to control (%)
100
80
60
40
20
0
Ctrl
CtIP
PIF1 siRNA efficency
cDNA relative to control (%)
Ctrl
PIF1
POLD3 siRNA efficency
cDNA relative to control (%)
Ctrl
POLD3
B
CTRL siRNA -DSB
CTRL siRNA +DSB
CtIP siRNA -DSB
CtIP siRNA +DSB
% ssDNA at DSB-1
25
20
15
10
5
0
200bp
C
-DSB
+DSB
EdU IP (% input)
5
4
3
2
1
0
Ori
DSB-2
DSB-4
D
n=3
Ctrl siRNA
SETX siRNA
Repair synthesis at DSB-2
(EdU-IP ratio +DSB/-DSB normalized to ori)
2
1,8
1,6
1,4
1,2
1
0,8
0,6
0,4
0,2
0
-3000
-1000
1000
3000
5000
DSB
distance from DSB (bp)

**Supplemental Table 1**: siRNA sequences used in this study

| | |
|---|---|
| BLM | AGCAGCGAUGUGAUUUGCAtt. |
| SETX | GAGAGAAUUAUUGCGUACUtt |
| CtIP | GCUAAAACAGGAACGAAUCtt |
| POLD3 | Silencer® Select validated siRNA Ambion s21045 |
| PIF1 | CCCUUCAGAGCCUAACCAAtt |

**Supplemental Table 2:** Primer sequences used in this study

| | Location of the genomic feature (hg19) | FW | REV | distance to DSB | application | related Figure(s) |
|---|---|---|---|---|---|---|
| **DSB-1** | chr22:38864102-38864108 | CCGCCAGAAAGTTTCCTAGA | CTCACCCTTGCAGCACTTG | 80bp | Cleavage assay, ChIP | Fig.S2B,Fig.S2D, Fig.3H-I, Fig.2E |
| | | ACCATGAACGTGTTCCGAAT | GAGCTCCGCAAAGTTTCAAG | 130bp | Resection | Fig.3F, Fig.4E |
| | | GGGTATGGAGCTGCCTCTAA | GACAAAGATGGCTGGAGGAG | 756bp | ChIP | Fig.S2B |
| | | ACAGATCCAGAGCCACGAAA | CCCACTCTCAGCCTTCTCAG | 850bp | Resection | Fig.3F, Fig.4E |
| **DSB-2** | chr9:130693171-130693177 | TCAAGTCTCAGGGACAAGCC | CTCCCGGGACGATTTCTCG | -2392bp | EdU-IP | Fig.S3D |
| | | GTGGGGGTCCCTTTTCAACC | AAGGCAGAGAGGGAGGAACA | -1321bp | EdU-IP | Fig.S3D |
| | | GCAGTCAGCACCCGAATAGAG | CATACCTACCACGAAGAAGCTGTT | -637bp | EdU-IP | Fig.S3D |
| | | ATGCTTTTCATAGCCGCTGAC | GGCGGATATCCCTCAACACTTC | -341bp | EdU-IP | Fig.S3D |
| | | TCCTCTCTCGGGTCCGC | GGAATGCGGCCAAAGCC | -100bp | EdU-IP | Fig.S3C-D, Fig.5C-D |
| | | AGACCTCGGTCCGGCT | GCAATGGGGATTTCACGCC | 35bp | EdU-IP | Fig.S3D |
| | | CCTACACTTAACCACTGAGCCG | GAGTCTGGCAAGGTGACGAAA | 479bp | EdU-IP | Fig.S3D |
| | | CTGTGCCGGATGAGTGTCTATG | ACACACCCATACTCACAGTACCT | 841bp | EdU-IP | Fig.S3D |
| | | GAGGAAGCCATCTGTGACTTAGG | GGACCCTGAGCTTTATCAGTTCTC | 2184bp | EdU-IP | Fig.S3D |
| | | GGGTGACAAGAGGGTGACTG | CCGTGGGGTTTTCCTGTTCA | 5388bp | EdU-IP | Fig.S3D |
| **DSB-3** | chr20:30946313-30946319 | CCTAGCTGAGGTCGGTGCTA | GAAGAGTGAGGAGGGGGAGT | 184bp | ChIP | Fig.3H |
| **DSB-4** | chr17:57184297-57184303 | ATCTCTTCAGGTCCAGCGCC | CCACCGTTCGCCCTACTTCT | 240bp | EdU-IP | Fig.S3C, Fig.5C-D |
| **ORI** | chr1:49240607-49244188 | TCAGACCCGAGCAATCACAG | CACCCCTATCCCGCATTCAC | | EdU-IP | Fig.S3C-D, Fig.5C-D |
| **Actin (ACTB)** | chr7:5566779-5569294 | AGCCGGGCTCTTGCCAAT | AGTTAGCGCCCAAAGGACCA | | ChIP | Fig.2E |
| **P0 (RPLP0)** | chr12:120634503-120639014 | GGCGACCTGGAAGTCCAACT | CCATCAGCACCACAGCCTTC | | RT-qPCR | Fig. 3E, Fig. S3A |
| **BLM** | chr15:91260579-91358686 | CCAGTGGTTCCAAGGCAAAG | GCTCCTGATGTCGCTTGAGAA | | RT-qPCR | Fig. 3E |
| **CtIP (RBBP8)** | chr18:20513295-20606449 | ACCCCCATGTCCGATACATA | TAGCTGCTTGTTCCATGTGC | | RT-qPCR | Fig. S3A |
| **POLD3** | chr11:74303575-74354105 | ACCAACAAGGAAACGAAAACAGA | GG TTCCGTGACAGACACTGTA | | RT-qPCR | Fig. S3A |
| **PIF1** | chr15:65107833-65113480 | GGGCGAGATGGGATTGTG | GCCCCGAGACACCGATAAGT | | RT-qPCR | Fig. S3A |
